# MKLP2 functions in early mitosis to ensure proper chromosome congression

**DOI:** 10.1101/2021.09.12.459884

**Authors:** Morgan S. Schrock, Luke Scarberry, Benjamin R. Stromberg, Claire Sears, Adrian E. Torres, David Tallman, Lucas Krupinski, Arnab Chakravarti, Matthew K. Summers

## Abstract

Mitotic kinesin-like protein 2 (MKLP2) is a motor protein with a well-established function in promoting cytokinesis. However, our results with siRNAs targeting MKLP2 and small molecule inhibitors of MKLP2 (MKLP2i) suggested a function earlier in mitosis, prior to anaphase. In this study we provide direct evidence that MKLP2 facilitates chromosome congression in prometaphase. We employed live imaging to observe HeLa cells with fluorescently tagged histones treated with MKLP2i and discovered a pronounced chromosome congression defect. We show that MKLP2 is essential for pole-based error correction as inhibited cells had a significant increase in unstable kinetochore-microtubule attachments due to impaired error correction of syntelic attachments. We propose that MKLP2 mediates kinetochore microtubule attachment stability by regulating Aurora Kinase activity and a downstream target, pHEC1 (Ser 55). Lastly, we show that MKLP2 inhibition results in aneuploidy, confirming that MKLP2 safeguards cells against chromosomal instability.

**Summary:** Schrock et al. demonstrate that the mitotic kinesin, MKLP2, is required for congression of chromosomes located near the spindle poles. They show that MKLP2 inhibition causes elevated active Aurora Kinase A, unstable microtubule kinetochore attachments, and impaired syntelic error correction.

## Introduction

Organisms require accurate and reliable segregation of their genetic material to grow and replace damaged cells. This process, known as mitosis, is a well-regulated, multi-step event with checkpoints in place to ensure accurate chromosome distribution to daughter cells. However, errors can occur, particularly in cancer, and may lead to daughter cells which inherit gains or losses of whole chromosomes, a cellular feature known as aneuploidy (Ben-David and Amon, 2020). Mitotic kinesin-like protein 2 (MKLP2) is a motor protein that harnesses energy generated through ATP hydrolysis to travel along dynamic microtubules during mitosis. MKLP2 is overexpressed in multiple cancers and was one of the highest ranking genes to associate with functional aneuploidy in cancer cells across 12 different cancer data sets (Carter et al., 2006). However, this association cannot be explained by our current understanding of MKLP2. At present, there is a well-established body of evidence to indicate that MKLP2 is required at the onset of anaphase to promote cytokinesis. Specifically, MKLP2 interacts with the chromosomal passenger complex (CPC, composed of: AURORA B, INCENP, SURVIVIN, BOREALIN) at the beginning of anaphase to maintain this heterotetrameric complex at the central spindle while homologous chromosomes segregate to opposite ends of the cell via bipolar microtubule attachments (Landino et al., 2017, Kitagawa et al., 2014, Hümmer and Mayer, 2009, Adriaans et al., 2020). Loss of MKLP2 allows formation of the ingression furrow between daughter cells but inhibits its completion, resulting in one binucleated cell instead of two daughter cells. Binucleation could explain MKLP2 association with polyploidy, multiple copies of the entire haploid genome, but does not explain the association with aneuploidy.

We had observed that inhibition of MKLP2 with paprotrain (Tcherniuk et al., 2010), a small molecule uncompetitive ATP inhibitor of MKLP2 (referred to here as MKLP2i^1^), caused a significant arrest in prometaphase characterized by the presence of chromosomes near the poles. Therefore the purpose of this study was to determine the role of MKLP2 prior to anaphase, given that the ‘pseudometaphase’ phenotype we observed could be caused by a chromosome congression defect or cohesion fatigue, both of which could contribute to chromosomal instability. We report here that MKLP2 functions in early mitosis to promote chromosome congression via correction of syntelic attachments and regulation of Aurora Kinase A (AURKA) and B (AURKB). Our experiments treating cells with second (MKLP2i^2^) and third (MKLP2i^3^) generation MKLP2 small molecule inhibitors (Ferrero et al., 2019, Labrière et al., 2016) revealed lagging chromosomes, chromosome bridges, and daughter cells with significantly distinct chromosome numbers, confirming that MKLP2 safeguards cells against aneuploidy.

## Results and Discussion

### MKLP2 deficiency leads to prolonged mitosis

In order to understand whether loss of MKLP2 has an effect prior to the onset of anaphase, we transfected HeLa cells with siRNAs targeting MKLP2 (siMKLP2) and quantified the metaphase index after release from double thymidine block (DTB) (Fig 1A). HeLa cells transfected with siMKLP2 exhibited reductions in MKLP2 (Fig 1B) and elevated metaphase index that remained significantly high four hours after cells began entering mitosis when the index of siCtrl-transfected had declined, indicating MKLP2-deficient cells feature prolonged metaphase. To validate this finding, we tracked individual HeLa H2B-GFP cells as they entered and exited mitosis. Treatment with MKLP2i^1^ (not shown) and MKLP2i^2^, revealed a dose-dependent increase in duration of mitosis such that 11 µM MKLP2i^2^ severely arrested cells (average 800 minutes) in metaphase (Fig 1C). While tracking cells, we also observed a dose-dependent decrease in the percentage of cells undergoing normal division (Fig 1D, 1E). The highest incidence of binucleation occurred at 3.7 µM MKLP2i^2^, while cells treated with 11 µM MKLP2i^2^ exhibited significantly more abnormal divisions (mitosis leading to 3 daughter cells or mitosis where cells exit mitosis with no apparent segregation of DNA). To confirm the mitotic arrest was specific to MKLP2, we performed an experiment in HeLa cells synchronized with DTB where MKLP2 siRNAs were transfected or small molecule MKLP2 inhibitors were added in parallel prior to mitosis. We confirmed that all modes of perturbing MKLP2 function increase the mitotic index albeit in a dose-dependent manner (Fig 1F) and found that MKLP2i^3^ (Pouletty, 2019) is ∼300 to 1,000 fold more potent than MKLP2i^2^ and MKLP2i^1^, respectively. Therefore we use this drug in all subsequent experiments.

**Figure 1.**
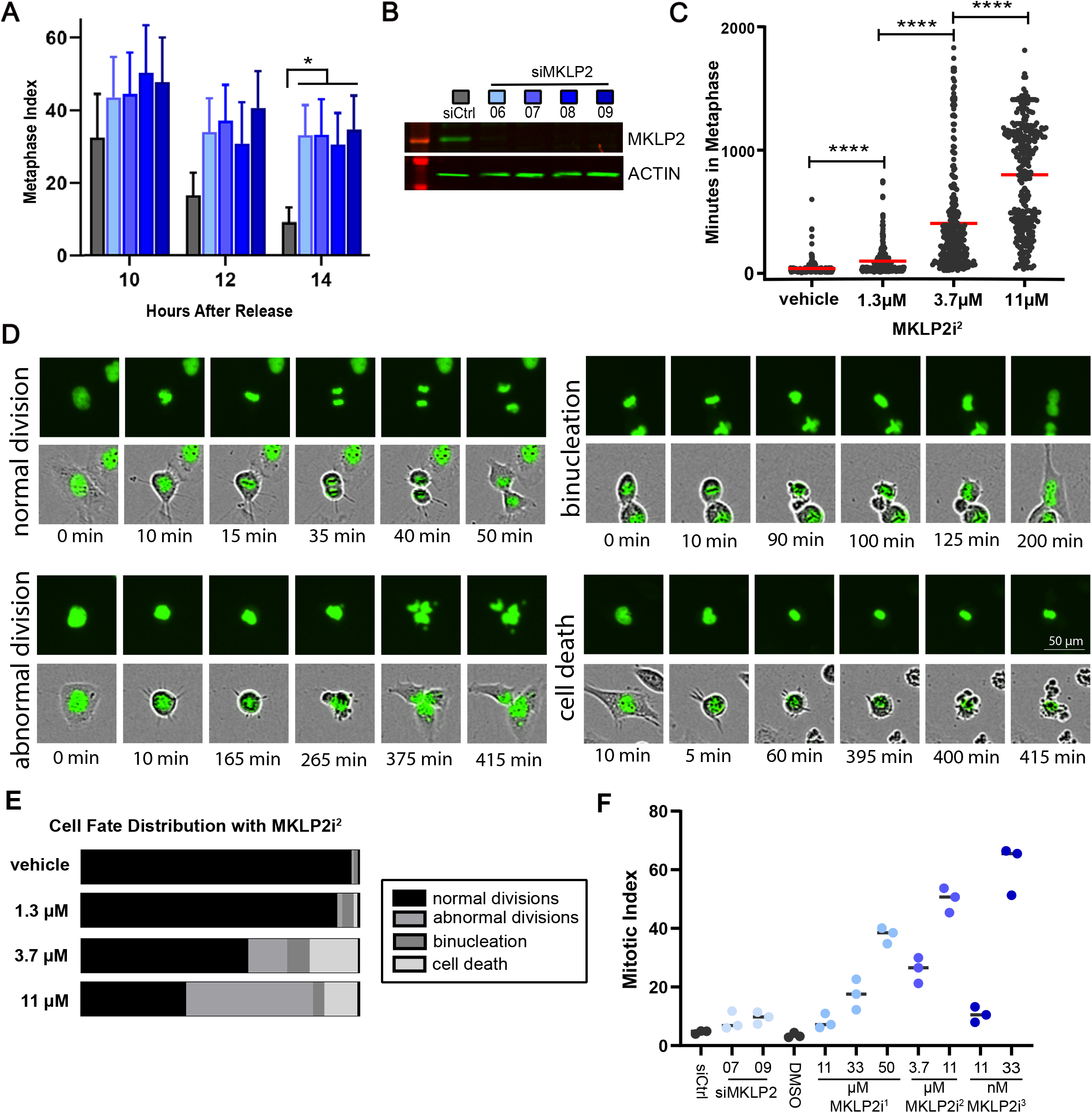
MKLP2 deficiency leads to prolonged mitosis. A) Metaphase index in HeLa cells transfected with the indicated siRNAs after release from DTB (ANOVA, * p<0.05, n>200 cells/condition, 2 independent experiments). B) Western blot of representative MKLP2 knockdown in cells treated as in A. C) Dose-dependent effect of MKLP2i^2^ on duration of metaphase (unpaired t test, ****p<0.0001, n>200 cells/condition, 3 independent experiments). D) Images portraying cell fates following treatment with MKLP2i^2^ for analysis quantified in C and E. Top panel-GFP channel, bottom panel-phase/GFP merge. E) Percentage of cells from C with indicated fates at the corresponding MKLP2i^2^ treatment dose. F) Metaphase index in HeLa cells 14 hours after release from DTB in the presence of various doses of MKLP2i^1^, MKLP2i^2^, MKLP2i^3^, and siMKLP2 (n= 50 cells/experiment, 3 independent experiments).

### MKLP2 deficiency leads to chromosome misalignment in prometaphase

We next examined chromosome movement under high magnification in live HeLa mCherry-H2B cells synchronized in mitosis with DTB. Treatment with MKLP2i^3^, resulted in a variety of abnormal mitotic phenotypes including chromosome bridges, lagging chromosomes, and a chromosome congression defect where the majority of cells properly congress to the metaphase plate but a small proportion of chromosomes remain at the poles (Fig 2A, supplemental movies). We refer to the congression defect as pseudometaphase when it results in a metaphase-like arrest greater than 100 minutes. Cells exhibited a dose-dependent increase in the manifestation of pseudometaphase (Fig 2B). To further confirm that these phenotypes were associated with inhibition of MKLP2 and not an off-target effect of MKLP2i^3^, we generated MKLP2 mutants with point mutations within the Switch II region (E413A) or P-loop region (G165E) of the motor domain that were previously shown to inhibit kinesin motor function (Browning et al., 2003). HeLa cells were co-transfected with siMKLP2 and an H2B-GFP plasmid to track transfected cells plus wild type MKLP2 or either of the rigor mutants plus. As expected, we observed the pseudometaphase phenotype in cells transfected with MKLP2 G165E mutant (13%) and MKLP2 E413A mutant (24%) compared to cells transfected with wildtype MKLP2 which exhibited no pseudometaphase cells (Fig 2C). To further confirm the presence of a chromosome congression defect in MKLP2 deficient cells, we quantified uncongressed chromosomes by determining the centromere distribution ratio (Stumpff et al., 2012). HeLa cells were synchronized in G2 with RO-3306, a CDK1 inhibitor, then released into DMSO or 33nM MKLP2i^3^ and fixed 30 min after release. We observed the pseudometaphase phenotype in MKLP2i^3^ treated cells (Fig 2E) and found a significant increase (p<0.001) in the centromere distribution ratios of MKLP2 deficient cells compared to vehicle treated cells (Fig 2F), confirming MKLP2 inhibition impairs chromosome congression.

**Figure 2.**
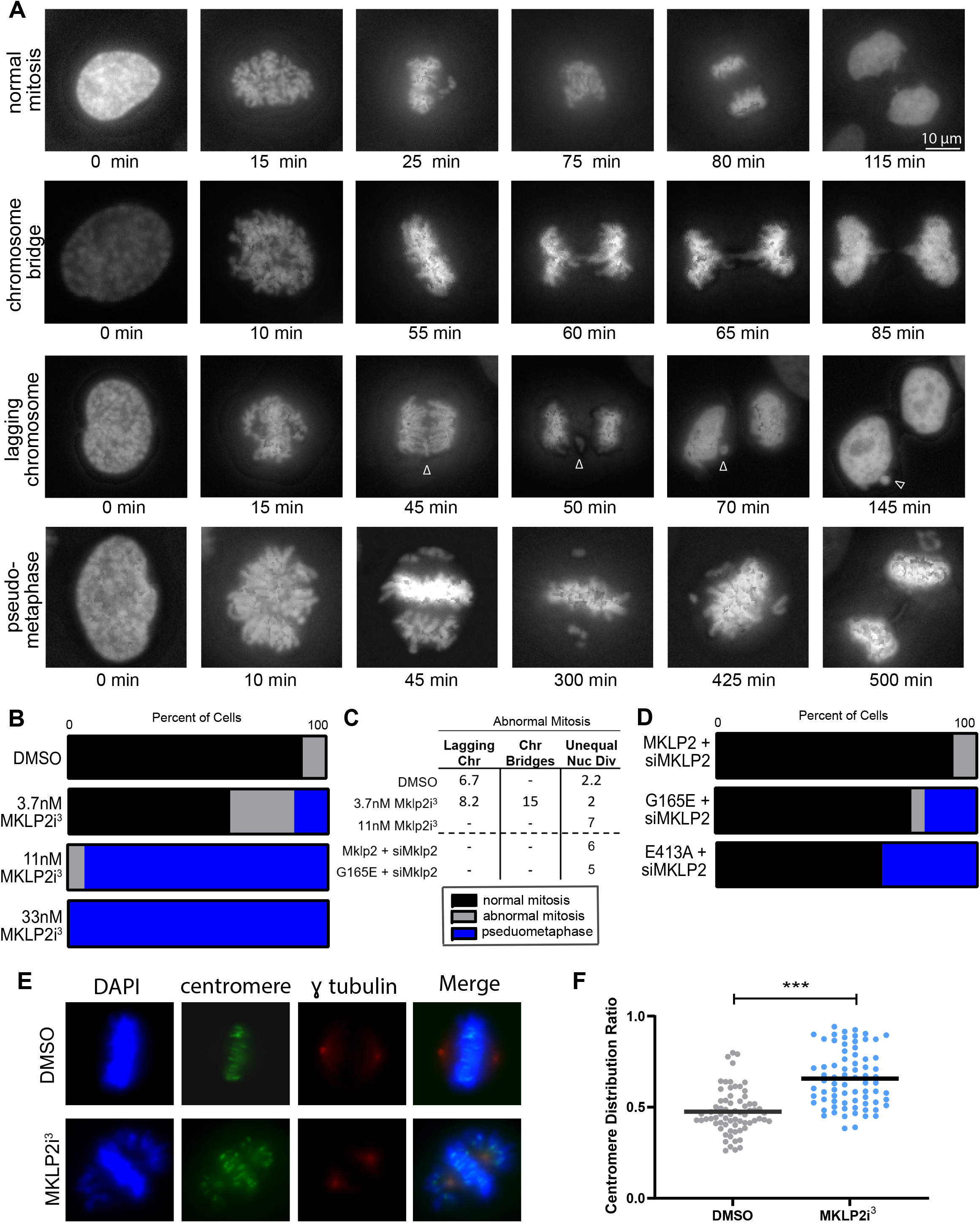
MKLP2 deficiency leads to chromosome misalignment in prometaphase. A) Images depicting chromosome segregation events captured in HeLa H2B mCherry cells following treatment with MKLP2i^3^ or DMSO. Arrow denotes lagging chromosome. B) Quantification of live imaging experiments from A indicating percentage of cells with the following outcomes: normal mitosis, abnormal mitosis, or pseudometaphase (prometaphase arrest >100 minutes from NEBD). (n>50 cells/ condition) C) Table reporting breakdown of abnormal events within cells that had abnormal mitosis. Numbers indicate percentage of cells with specified fate out of total cells. D) Live imaging quantification of mitotic outcomes as in B in HeLa mCherry H2B cells transfected with siMKLP2 + one of the following: vector, TAP-KIF20A (WT), TAP-KIF20A G165E (motor mutant), TAP-KIF20A E413A (rigor motor mutant). (n>50 cells/ condition) D) IF of γ-tubulin and centromere proteins in HeLa cells synchronized at G2/M with RO-3306 then released into DMSO or 33nM MKLP2i^3^ for 30 minutes. E) Quantification of centromere distribution ratio in HeLa cells treated with 33nM MKLP2i^3^ (unpaired t test, p<0.001, n = 100, 3 independent experiments).

### MKLP2 inhibition results in unstable kinetochore-microtubule attachments and impaired syntelic error correction

We hypothesized that misaligned chromosomes in the pseudometaphase phenotype were unable to reach the cell equator due to an inability to form stable kinetochore-microtubule (KT-MT) attachments. To test this idea we employed the cold depolymerization assay to identify stable microtubules (DeLuca et al., 2016). Following G2 synchronization with RO-3306, we released cells into DMSO or 33 nM MKLP2i^3^ then incubated cells with cold media. We then measured the intensity of the remaining cold-stable microtubules and found that mitotic cells in the presence of 33 nM MKLP2i^3^ exhibited significantly less (p<0.0001) tubulin compared to DMSO treated cells (Fig 3A, 3B). To determine whether unstable KT-MT attachments in MKLP2i^3^ treated cells are due to issues in correction of misaligned chromosomes, we employed the monastrol washout assay where treatment with monastrol generates an abundance of syntelic attachments which are then allowed to correct in a washout period. The cells do not exit mitosis due to the presence of the proteasome inhibitor, MG132 (Lampson et al., 2004). We found that the majority of DMSO treated cells were able to create bipolar spindles with aligned chromosomes after monastrol washout, while MKLP2i^3^ treated cells created bipolar spindles but with numerous misaligned chromosomes, resembling the pseudometaphase phenotype (Fig 3C, 3D). These experiments indicate that the unstable KT-MT attachments that characterize the pseudometaphase phenotype with MKLP2 inhibition are due to impairment of a syntelic error correction mechanism.

**Figure 3.**
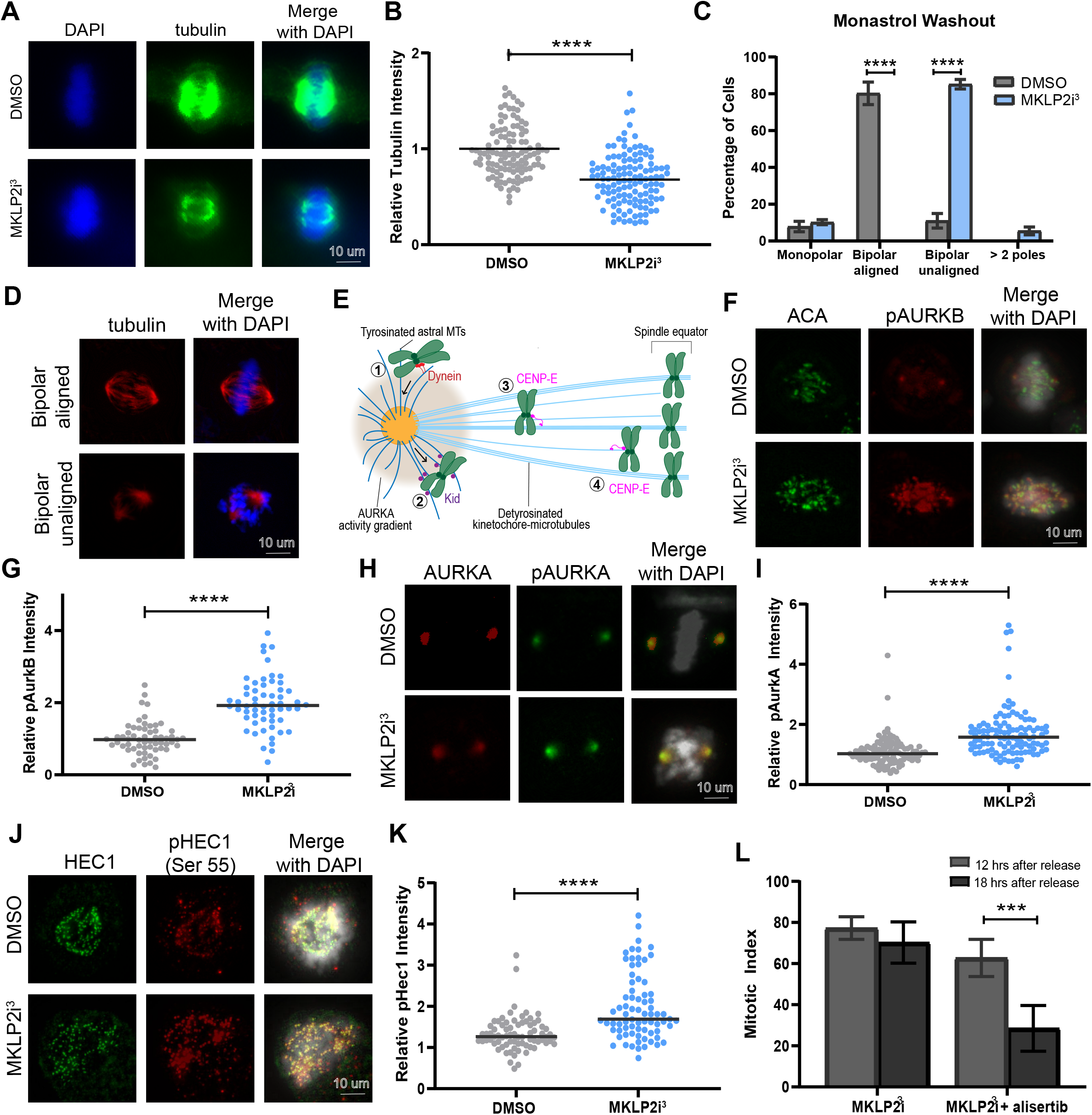
MKLP2 deficiency results in unstable microtubule kinetochore attachments, impaired syntelic error correction, and upregulation of Aurora Kinases. A) IF of cold-stable microtubules in Hela cells treated with DMSO or 33nM MKLP2i^3^. B) Quantification of relative tubulin intensity from cells in A (n = 100, 3 independent experiments unpaired t test, ****p<0.0001). C) Effect of 33nM MKLP2i^3^ on chromosome congression after monstraol washout (unpaired t test, ****p<0.0001, n = 100, 3 independent experiments). D) Representative images pertaining to C. E) schematic of pole-based error correction. (1) Chromosomes located outside the interpolar region are transported poleward along tyrosinated astral MTs via dynein. (2) Kid generates polar ejection forces that propel chromosomes away from the poles. While in proximity to the poles, an AURKA activity gradient phosphorylates pHEC1 and destabilizes KT-MT attachments. (3) Unattached chromosomes are transported by CENP-E via lateral attachments to detyrosinated MTs toward the spindle equator. (4) Once the chromosome is mono-oriented, CENP-E facilitates its movement via lateral attachment to k-fibres to the metaphase plate. F) Images of IF staining for pAURKB and anti-centromere antibody (ACA) in mitotically synchronized HeLa cells with and without 33nM MKLP2i^3^. G) Relative pAURKB levels after 33 nM MKLPi^3^ treatment pAURKB intensities were normalized to ACA. (unpaired t test, ****p<0.0001, n = 100, 3 independent experiments). H) Images of IF staining for pAURKA and total AURKA in mitotically synchronized HeLA cells with and without 33nM MKLP2i^3^. I) Graph depicting increased pAURKA (unpaired t test, ****p<0.0001) in cells treated with MKLP2i^3^. pAURKA intensities were normalized to AURKA. (n = 100, 3 independent experiments) J) Images of IF staining for pHEC1 and total HEC1 in mitotically synchronized HeLa cells with and without 33nM MKLP2i^3^. K) pHEC1 levels in cells treated with 33nM MKLP2i^3^. pHEC1 intensities were normalized to HEC1. (unpaired t test, ****p<0.0001); n = 100, 3 independent experiments). L) Mitotic index of HeLa cells synchronized with DTB and treated with 33nM MKLP2i^3^ or 33nM MKLP2i^3^ + 12.5nM alisertib (AURKA inhibitor). (unpaired t test, ***p<0.001, n>100 cells, 3 independent experiments).

### Aurora Kinases including AURKA, a key component of pole-based error correction, is upregulated in MKLP2 inhibited cells

The canonical error correction pathway is mediated by centromere-enriched AURKB-catalyzed phosphorylation of HEC1, a component of the NDC80 complex that serves as the main link between kinetochores and microtubules (Krenn and Musacchio, 2015, Lampson and Cheeseman, 2011). AURKB phosphorylation of HEC1 N-terminal “tail” (including S55, pHEC1 hereafter) domain causes destabilization of mal-oriented KT-MT attachments thus providing a fresh opportunity for microtubules to make correct amphitelic attachments, where each sister kinetochore is attached to microtubules emanating from opposite poles (Wimbish and DeLuca, 2020). The application of force from opposing poles generates tension which then stabilizes correct KT-MT attachments. A distinct but complementary pathway known as pole-based error correction, summarized in Fig 3E, promotes proper alignment of chromosomes initially located near the pole, which are more likely to form syntelic attachments, via microtubule motor-based transportation of chromosomes to the cell equator (Ye et al., 2015, Chmátal et al., 2015, DeLuca, 2017). Briefly, chromosomes located outside the interpolar region are transported poleward via dynein where the chromokinesin kid generates polar ejection forces that propel chromosomes away from the poles. Near the poles, AURKA phosphorylates substrates, including HEC1 S55, which destabilizes KT-MT attachments. Mono-oriented or unattached chromosomes are then transported by CENP-E toward the spindle equator where they become bioriented (Kapoor et al., 2006). This pole-based error correction mechanism helps ensure congression of chromosomes that are initially located near the poles at nuclear envelop breakdown and prevents their entrapment near the poles.

In order to understand how MKLP2 inhibition regulates KT-MT attachment and error correction, we sought to determine the relative levels of active AURKB, AURKA, and the downstream target pHEC1. Immunofluorescence (IF) analysis of phosphorylated AURKB and AURKA revealed significant increases (p<0.0001) in the active states of both kinases in MKLP2i^3^ treated cells (Figs 3F-3I). Because AURKA activity has been shown to help align pole-based chromosomes via error correction through phosphorylation of HEC1 (Ye et al., 2015), we examined the contribution of MKLP2 inhibition to pHEC1 using a phospho-specific antibody against pS55 in HeLa cells. Compared to control cells, MKLP2i^3^ treatment caused a significant increase in pHec1 staining (Fig 3J, 3K; p<0.0001). In an attempt to tease out which Aurora Kinase was responsible for impaired error correction in MKLP2i^3^ treated cells, we synchronized HeLa cells with a DTB and treated with MKLP2i^3^ in combination with alisertib, an AURKA inhibitor. As expected, 33 nM MKLP2i^3^ caused a pseudometaphase arrest in the majority of cells, however, treatment with a low dose (12.5 nM) of alisertib rescued this arrest six hours later (Fig 3L) suggesting that elevated AURKA is responsible for the congression defect observed in MKLP2 inhibited cells. However, we cannot conclude whether increased pAURKB destabilizing KT-MT attachments contributes to impaired error correction or is simply a consequence of the persistence of syntelic attachments due to impaired pole-based error correction. We performed a similar rescue experiment with the AURKB inhibitor, barasertib (not shown), that did not show rescue of the mitotic delay caused by MKLP2i. However, AURKB inhibition in itself induces a moderate mitotic arrest (Gurden et al., 2016), confounding the interpretation. While more work is needed to more completely characterize the contributions of AURKB in the MKLP2i-induced congression defect, our data suggests that under normal conditions MKLP2 negatively regulates AURKA to promote syntelic error correction.

### MKLP2 deficiency results in aneuploidy and contributes to chromosomal instability

To determine whether congression defects caused by MKLP2 deficiency contribute to cellular aneuploidy, we employed satellite enumeration probes to quantify individual chromosomes via FISH in HeLa cells. We found that a 48 hour treatment of 33 nM MKLP2i^3^ significantly increased the frequency at which cells carried a chromosome number distinct from the mode (chromosomes 2, 7, and 15, Fig 4A, B). This data suggests that MKLP2 association with functional aneuploidy observed by Carter et al. (2006) is due to MKLP2 expression limiting chromosomal instability (CIN), the gain or loss of chromosomes at a higher incidence than normal. To further evaluate the role of MKLP2 in cells exhibiting CIN, we evaluated colorectal cancer (CRC) cell lines to compare cell lines with CIN (microsatellite stable, MSS) to cells without CIN (microsatellite unstable, MSI). Utilizing publicly available data (DepMap, Broad (2019)), we evaluated the effect of MKLP2 knockdown via RNAi or loss by CRISPR and found that cell lines exhibiting CIN (MSS) were significantly more sensitive to loss of MKLP2 compared to CRC cell lines without CIN (Fig 4C, D), suggesting that cancer cell lines with ongoing CIN require MKLP2 to limit excessive CIN.

**Figure 4.**
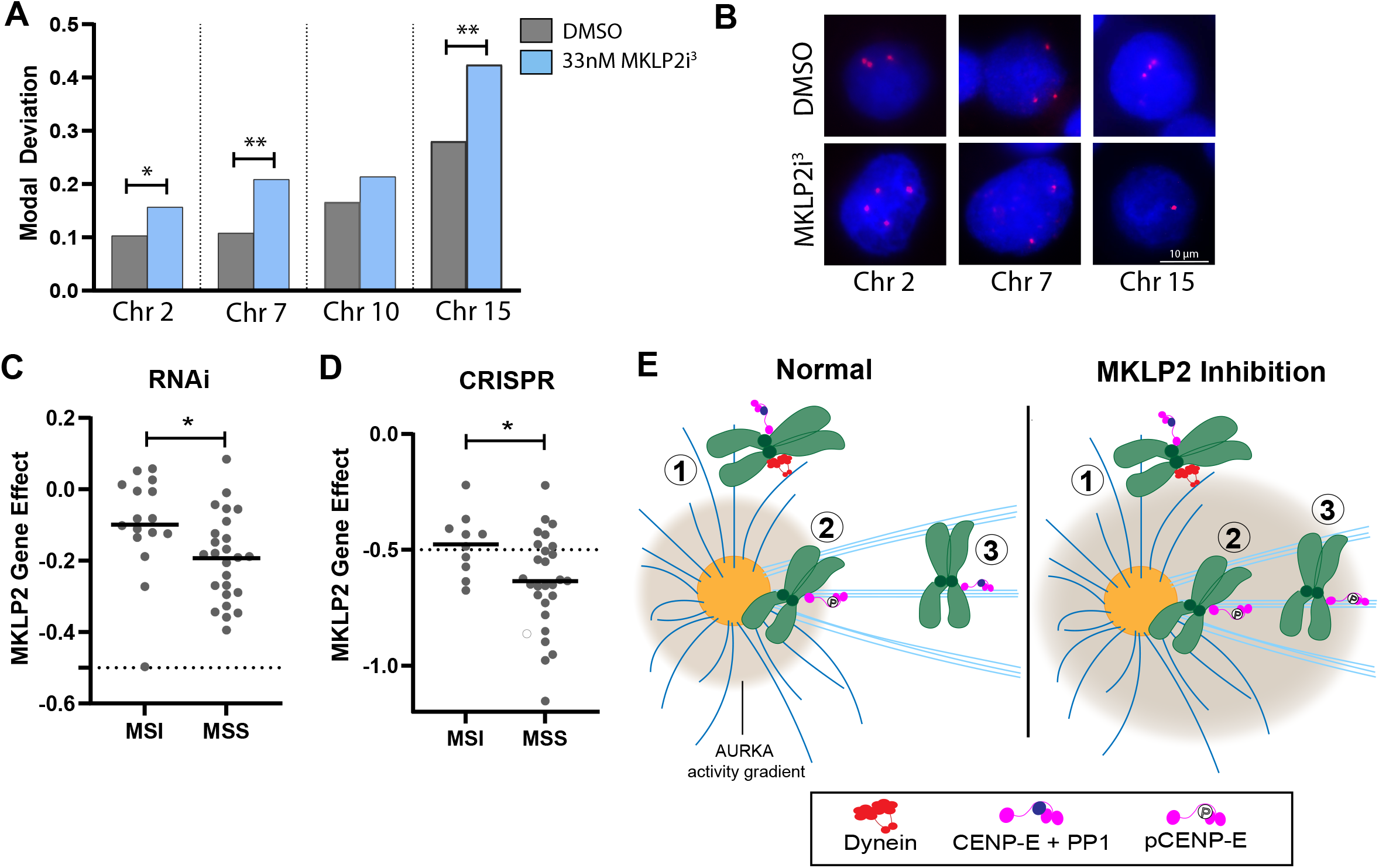
MKLP2 loss causes aneuploidy and exacerbates CIN. A) Fraction of cells with chromosome number deviating from the mode (3 chromosomes/cell) for 4 chromosomes (2, 7, 10, and 15). (Chi square test, *p<0.05, **p<0.01, n > 200 cells/condition, 2 independent experiments) B) Images of chromosome enumeration probes (red) from A. C) Gene effect scores of CRC cell lines with CIN (MSS, n=25) and without CIN (MSI, n=16) when MKLP2 was depleted via RNAi (unpaired t test, * p<0.05). D) Gene effect scores of CRC cell lines with CIN (MSS, n=27) and without CIN (MSI, n=10) when MKLP2 was depleted via CRISPR (unpaired t test, * p<0.05). E) Proposed model of the consequences of MKLP2 inhibition on chromosome alignment. Under normal conditions (1) Chromosomes outside the interpolar region contain PP1 bound CENP-E and are motored toward the poles via dynein. (2) Once within the AURKA activity gradient near the pole, AURKA phos CENP-E at Thr422, displacing PP1 and destabilizing KT-MT attachments. (3) CENP-E motors laterally attached chromosomes towards the cell equator and once outside the AURKA activity gradient, PP1 binds CENP-E Thr422 where it dephosphorylates CENP-E, Ndc80 and KNL1 enabling the conversion of lateral to end-on KT-MT attachments. However in cells with impaired MKLP2 motor activity, the increased AURKA activity gradient prevents PP1 binding to CENP-E at Thr422 thus preventing the conversion of unstable KT-MT attachments to stable end-on attachments.

## Conclusions

In this manuscript we report that MKLP2 functions in early mitosis to facilitate congression of pole-based chromosomes via syntelic error correction. Given that the early mitotic phenotype becomes more pronounced with increasing concentration of MKLP2 inhibition, whereas binucleation is apparent at lower doses (Figs 1C and 2B), we propose that the role of MKLP2 in early mitosis requires a smaller amount of functional enzyme than is required for its role in cytokinesis. As such, RNAi may not deplete sufficient amounts of the protein to reveal the phenotype consistently. In support of this idea, we evaluated the consequence of MKLP2 loss via RNAi compared to CRISPR using Project Achilles data via the DepMap Portal. We found that partial loss of MKLP2 function via RNAi is generally tolerated, while total loss of function achieved via CRISPR was not, suggesting a small amount of MKLP2 is sufficient for viability (Suppl Fig 1A).

While our data clearly shows a drastic inability of cells to correct syntelic attachments in the presence of MKLP2 inhibition (Fig 3C), IF data (Fig 3H, 3I) and an alisertib rescue experiment (Fig 3L) suggest that elevated AURKA activity is responsible. This raises the question as to how that can be when studies report an AURKA activity gradient is integral to the correction of syntelic attachments (Ye et al., 2015, Chmátal et al., 2015). Previous analyses shed light on this question by demonstrating antagonistic regulation of CENP-E by the Ser/Thr phosphatase protein phosphatase 1 (PP1) and AURKA (Kim et al., 2010). In this model, summarized in Fig 4E, PP1 is bound to CENP-E on pole localized chromosomes before dynein motors move them towards the pole. Once chromosomes are within the AURKA activity gradient, AURKA phosphorylates CENP-E displacing PP1 and destabliziling KT-MT attachments. CENP-E motors move laterally attached chromosomes towards the cell center and once outside the AURKA activity gradient, PP1 can bind CENP-E once again where it dephosphorylates not only CENP-E but also Ndc80 and KNL1, enabling the conversion of lateral KT-MT attachments to stable end-on attachments (Kim et al., 2010). Given this PP1/AURKA phosphorylation switch, we propose that excessive AURKA activity caused by inhibition of MKLP2 prevents PP1 binding to CENP-E and the dephosphorylation cascade necessary to create stable KT-MT attachments (Fig 4E). Indeed, preventing PP1 rebinding to CENP-E inhibits the formation of stable kinetochore attachments to microtubules (Kim et al., 2010).

How exactly MKLP2 negatively regulates AURKA activity is still an unanswered question. The fact that the G165E and E413A MKLP2 rigor mutants recapitulate the pseudometaphase phenotype (Fig 2D) not only confirms the specificity of our finding to MKLP2, but also suggests that the motor activity of MKLP2 is important to syntelic error correction and regulating AURKA activity. MKLP2 has processive plus-end directed motor activity (Adriaans et al., 2020), suggesting that under normal conditions MKLP2 motors substrates towards the cell equator to negatively regulate AURKA activity. In support of this idea, MKLP2 rigor mutants concentrate at the spindle poles and MKLP2i^1^ has a similar impact (Adriaans et al., 2020, Krupina et al., 2016). Interestingly, AURKA interacts with and is activated by the CPC (Katayama et al., 2008), similarly to AURKB. The AURKA-CPC complex is thought to mediate phosphorylation of AURKA substrates at the kinetochore/centromere throughout mitosis (DeLuca et al., 2018). Little is known about the regulation of this complex, but it appears that while it does not require microtubules to form, proper localization does require microtubules (DeLuca et al., 2018). Thus, an attractive possibility is that MKLP2 regulates AURKA in a fashion analogous to AURKB through transportation of the AURKA-CPC complex where loss of MKLP2 function leads to an accumulation of active AURKA at the poles (Adriaans et al., 2020, Serena et al., 2020). Consistent with our hypothesis that only a small fraction of MKLP2 is required for this function, it has been difficult to visualize this pool of AURKA-CPC at endogenous levels (DeLuca et al., 2018). Alternatively, evidence suggests that MKLP2 interacts with BRCA1, which negatively regulates AURKA activity (Ertych et al., 2016, Hill et al., 2014). Further work is required to determine how these mechanisms may contribute to regulation of AURKA activity by MKLP2.

## Supporting information

Suppl movie 1. DMSO

Suppl movie 2. 33nM MKLP2i3

Suppl movie 3. 3.7nM MKLP2i3

Suppl movie 4. siMKLP2 + MKLP2

Suppl movie 5. siMKLP2 + G165E

Suppl movie 6. siMKLP2 + E413A

## Acknowledgements

This work was supported by an American Brain Tumor Association Basic Research Fellowship sponsored by an Anonymous Corporate Partner (MSS), GM108743 (MKS), GM112895 (MKS), R01CA227874-03 (AC), R01CA1145128-06 (AC), R01NS104332 (AC), R01CA227874 (AC), R01CA188228-06 (AC), and The Ohio State University Comprehensive Cancer Center (OSU CCC)/Department of Radiation Oncology start-up funds (MKS). Live cell imaging experiments were performed at the OSU Neuroscience Imaging Core supported by P30 NS104177. Research reported in the publication was supported by The Ohio State University Comprehensive Cancer Center and the NIH under grant number P30 CA016058. The authors thank OSU CCC’s Genomics Shared Resource for technical support and Medicinal Chemistry Shared Resource, an NCI subsidized Shared Resource supported by NCI-CCSG P30CA016058, for synthesis of MKLP2i^2^ and MKLP2i^3^. This project was also supported by Award Number UL1TR002733 from the National Center for Advancing Translational Sciences. The content is solely the responsibility of the authors and does not necessarily represent the official views of the National Center for Advancing Translational Sciences or the NIH. All authors declare no competing financial interests. We thank Dr. Katharine Ullman for gifting us the HeLa mCherry H2B GFP tubulin cells. MSS particularly thanks Paula Monsma for providing essential training and advice for live cell imaging experiments and Jennifer Morse, for critical expertise with the Cytocell satellite enumeration probe experiments.

## Author contributions

M.S.S. performed all experiments, L.S., B.R.S., and A.E.T. assisted in conducting experiments. C.S. analyzed cells in Fig 1D. L.K. analyzed Fig 4 experiments. D.T. wrote the code for generating SME projections and compiling into a movie. M.S.S. and M.K.S. designed and analyzed the experiments and wrote the manuscript. A.C. and M.K.S provided project supervision and support. All authors reviewed and approved the manuscript.

## Figure titles and legends

**Supplemental Figure 1**. A) Graph depicting viability dependency score among cancer cell lines with depletion of MKLP2 via RNAi or CRISPR (n=437, unpaired t test; ****p<0.0001).

**Supplemental movie 1**. Epifluorescent images taken every 5 minutes of HeLa mCherry H2B GFP tubulin cells after DTB and treated with DMSO. Images were compiled into a time-lapse movie of SME projections.

**Supplemental movie 2**. Epifluorescent images taken every 5 minutes of HeLa mCherry H2B GFP tubulin cells after DTB and treated with 33nM MKLP2i^3^. Images were compiled into a time-lapse movie of SME projections.

**Supplemental movie 3**. Epifluorescent images taken every 5 minutes of HeLa mCherry H2B GFP tubulin cells after DTB and treated with 3.7nM MKLP2i^3^. Images were compiled into a time-lapse movie of SME projections.

**Supplemental movie 4**. Epifluorescent images taken every 5 minutes of HeLa mCherry H2B GFP tubulin cells after DTB and transfected with siMKLP2 + pCS2 TAP KIF20A. Images were compiled into a time-lapse movie of SME projections.

**Supplemental movie 5**. Epifluorescent images taken every 5 minutes of HeLa mCherry H2B GFP tubulin cells after DTB and transfected with siMKLP2 + pCS2 TAP KIF20A G165E. Images were compiled into a time-lapse movie of SME projections.

**Supplemental movie 6**. Epifluorescent images taken every 5 minutes of HeLa mCherry H2B GFP tubulin cells after DTB and transfected with siMKLP2 + pCS2 TAP KIF20A E413A. Images were compiled into a time-lapse movie of SME projections.

## Materials and Methods

### Lead Contact

Further information and requests for resources and reagents should be directed to and will be fulfilled by the Lead Contact, Matthew K. Summers.

## EXPERIMENTAL MODEL AND SUBJECT DETAILS

HeLa and HeLa mCherry H2B GFP Tubulin cells (female) were maintained in DMEM, 10% FBS, 1% Pen Strep except when conducting live imaging experiments when they were maintained in Leibovitz without phenol red, 10% FBS, 1% Pen Strep. Cell lines were verified with microsatellite genotyping by The Ohio State University Comprehensive Cancer Center Genomics Shared Resource and Mycoplasma tested negative with SIGMA Lookout Mycoplasma PCR Detection Kit.

## METHOD DETAILS

### Cell Synchronization

For experiments shown in Figs 2E-F, Fig 3A-B, Fig 3F-K cells were synchronized in G2 with a single thymidine block x 20 hrs followed by release into 5 µM CDK1 inhibitor, RO-3306, for 7 hours, then released into DMSO or drug. Experiments began when control cells were just entering metaphase-30 min after release from RO-3306. For experiments shown in Fig 1 (all panels), Fig 2A-D, Fig 3L cells were synchronized with a double thymidine block consisting of 2mM Thymidine x 18 hrs, followed by release in normal media x 6 hrs, followed by 2mM thymidine x18 hr, release in normal media. Cells entered mitosis approximately 10-12 hours after release. Cells were rinsed three times with PBS, which was warmed to improve synchronizations.

### Cloning

The pCS2 TAP MKLP2 plasmid was generated by obtaining MKLP2 cDNA from the Harvard PlasmID repository and ligating into the pCS2 TAP vector via gateway technology. To generate the G165E p-loop and E413A switch II MKLP2 motor mutants, primers noted in the Key Reagent Table were used with NEB Q5 Hot Start Master Mix to amplify pCS2 TAP MKLP2 plasmid. Mutations were confirmed with sequencing.

### Immunofluorescence

For immunofluorescence cells are seeded onto round coverslips in one well of a 24 well plate. When washing or adding reagents, instead of directly adding to the well, we found that moving coverslips into wells that had been prefilled with wash/primary/block etc. greatly minimized loss of mitotic cells. All immunofluorescence experiments were imaged with Invitrogen EVOS M7000 at 100x. Cells from Fig 3 (all panels) were imaged in a z stack.

For immunofluorescence detection of α CREST, α γ-tubulin, cells were fixed in 4% PFA for 20 min, then ice cold methanol stored at -20°C for 10 minutes, blocked in 20% goat serum at room temperature (RT) for one hour, then incubated with human α CREST (Antibodies Inc; 1:500 in AbDil (1% BSA, 0.1% Triton X-100, 0.1% NaAzide in TBS pH 7.4)) overnight at 4°C. After a wash, the second antibody, α γ-tubulin (SIGMA GTU-88; 1:1,000 in AbDil) was incubated for 1 hour at RT. Corresponding secondary antibodies were also diluted 1:500 in AbDil and incubated for one hour at RT. Cells were counterstained with Hoechst (SIGMA) 1:10,000 during a 5 min wash and then mounted with Fluoromount media. All washes were done three times for 5 min in TBS

For cold-induced microtubule stabilizing IF with α-tubulin, 30 min after HeLa cells synchronized in mitosis were released into DMSO or 50nM compound 38, media was replaced with ice-cold DMEM (containing either drug or DMSO) and placed on ice for 10 min. Cells were washed once with PHEM buffer (60mM PIPES, 25mM HEPES, 20mM EGTA, 8mM MGSO_4_, pH 7.0) for 5 minutes, then permeabilized in 0.5% TritonX-100 in PHEM buffer for 4 minutes at RT. Cells were fixed in 4% PFA for 20 min at RT, then blocked in 10% goat serum diluted in PHEM at RT for one hour. Rat α α-tubulin (clone YL1/2) was diluted 1:200 in 5% goat serum diluted in PHEM and incubated at RT for one hour. Corresponding secondary antibody was diluted 1:300 and incubated RT for 1 hour. Cells were counterstained with Hoechst (SIGMA) 1:10,000 during a 5 min wash. All washes were done three times for 5 min with PHEM. Relative spindle intensity was measured with by quantifying α-tubulin staining in FIJI and normalized to DAPI intensity.

For pHec1 Ser55 normalized to total Hec1, cells were pre-extracted with pre-warmed BRB80 (80mM PIPES pH 6.8, 5mM EGTA, 1mM MgCl_2_, 0.5% Triton X) for 2 minutes at 37°C. Cells were then fixed in 4% PFA for 15 minutes at room temperature, washed twice for 5 minutes in PBST (0.1% Triton X-100), and blocked in blocking buffer (3% BSA, 0.1% Triton X-100, PBS) for 30 minutes at room temperature. Primary antibodies (GTX pHec1 Ser55, GTX Hec1 9G3.23) were diluted 1:3,000 in blocking buffer and incubated overnight at 4°C. The next morning cells were washed twice and incubated with secondary antibodies (1:500 diluted in blocking buffer) for 1 hour at room temperature. Cells were washed three times in PBST, counterstained with Hoechst and mounted using fluoromount. Mitotic cells were imaged at 100x with EVOS Cell Imaging System (Thermo Fisher Scientific). Intensities were measured in FIJI by splitting channels, background subtracting, then encircling foci contained within the chromosomal area to specify area to be measured. Intensity of pHec1 was normalized to total Hec1 intensity in 3 biological replicates (25 cells/ condition).

For immunofluorescence of pAURKB, ACA (Figs 3F, 3G), pAURKA, and total AURKA (Figs 3H, 3I), cells were fixed in 4% PFA x 20 min at RT, then permeabilized in 0.5% Triton X-100 × 2 min. After a brief rinse in PBS, cells were blocked in 5% BSA (PBS) x 30 min, then incubated with primary overnight at 4°C (all primaries were prepared in 5% BSA at 1:1000 dilution). The next day coverslips were washed 3 times with PBS (5 min each) then secondary antibodies (1:500 diluted in 5% BSA) were incubated in the dark x 1 hr at RT then washed three times (5 min each) with PBS. Coverslips were mounted onto slides with Invitrogen ProLong Gold Anti-Fade with NucBlue mountant and allowed to cure in the dark ON.

### Live imaging

For live cell imaging shown in Fig 2A-2C, HeLa H2B mCherry GFP tubulin cells were synchronized with a double thymidine block and released into normal DMEM. For experiment shown in Fig 2B with various doses of MKLP2i^3^, media was replaced with Leibovitz media containing 33nM MKLP2i^3^ six hours after second release. For live imaging experiments with transfection of MKLP2 rigor mutants (Fig 2C-D), MKLP2 plasmids (G165E or E413A) were transfected one hour after release from first thymidine block with LT1 according to MIrus Bio protocol. Transfection mixture containing siMKLP2 and RNAiMax (prepared according to Thermo Fisher protocol) was overlayed onto cells after addition of media containing thymidine six hours after release from first thymidine block. Cells were imaged using-wide field imaging on Nikon TiE microscope and captured on an Andor Ultra EMCCD camera. Cells were imaged every 5 minutes for 10 hours using 60x or 20x and 1 µm step size (total steps= 14). 2D projections of z stacks were generated using smooth manifold extraction (Shihavuddin et al., 2017) and assembled into movies using a script that will be submitted as supplemental information.

For live imaging data shown in Fig 1C-E, cells were released from double thymidine block into various doses MKLP2i2 then imaged in a single plane of focus with IncuCyte ZOOM Live-Cell Analysis System at 20x every 5 minutes x 48 hours.

### FISH with Cytocell satellite enumeration probes

To determine aneuploidy following treatment with MKLP2i^3^, we performed Fluorescence in situ hybridization with Cytocell satellite enumeration probes (Oxford Gene Technology). To harvest, trypsinized cells were centrifuged (1,000 rpm) x 3 minutes and resuspended in ¼ ‘‘ supernatant. Pre-warmed (37°C) hypotonic 0.075M KCl (6mL) was added slowly to each sample, then samples were incubated in a 37°C water bath x 15 minutes. Ten drops of freshly prepared Carnoys solution (1:3 glacial acetic acid: methanol) was added to each sample, then centrifuged x 10 minutes. Supernatant was carefully aspirated leaving ¼ ‘‘ for resuspension, then 8 mLs Carnoys was added and cells were fixed x 30 min at 4°C. Samples were centrifuged x 3 minutes, supernatant aspirated leaving ¼ ‘‘ for cell resuspension. Cells (10ul Carnoy’s resuspension) were dropped onto clean slides warmed to 65°C on a hot plate. Cells dried to slide by remaining on hot plate x 15 seconds, then a pap marker was used to outline the dried suspension circumference on the underside of the slide. Cells were denatured by incubating in 2X SSC x 2 minutes followed by a brief protein digestion (1:20 proteinase K: 2X SSC) x 12 minutes at 37°C. Cells were incubated once more in 2X SSC x 2 minutes, followed by dehydration in an ethanol series (70%, 85%, 100%) x 2 minutes each. Slides were air dried by leaning up and resting on a kimwipe. Slides and probes mixture (3 µl probe + 7 µl Cytocell hybridization solution) were warmed in a 37°C incubator, then 10 µl probe/hybridization solution was overlaid onto each cell sample. A coverslip was applied and the edges were sealed with rubber cement. Slides were placed on a 75°C hot plate x 2 minutes, then incubated overnight at 37°C in the dark. The next day rubber cement was carefully and completely removed, then slides were incubated in 0.4X SSC x 2 minutes at 72°C. Next slides were immersed in 2X SSC, 0.05% Tween-20 × 30 seconds, followed by the ethanol series again (70%, 85%, 100%) for 2 minutes each. Slides were allowed to air dry on a kimwipe. Prior to applying the coverslip, 5 µl diluted Hoechst (1 µl Hoechst in 8 ml PBS) and 5 µl fluoromount (Southern Biotech) were applied to the cell area. Slides were allowed to dry at room temperature overnight in the dark. Images were captured with Invitrogen EVOS M7000.

## QUANTIFICATION AND STATISTICAL ANALYSIS

Image analysis was performed in ImageJ/FIJI. The merge of z stack images was used for analysis. For centromere distribution ratios in Figs 2E, 2F, cells were imaged in a plane of focus that captured both centrosomes using a Leica DM5500B fluorescent microscope as described previously (Stumpff et al., 2012). Centromere intensity per cell quarter was measured using FIJI. The centromere distribution ratio (r = γ_1_ + γ_2_ / ε) was then calculated for each cell by dividing the sum of the intensity of the outer quarters closest the poles by the sum of the intensity of the inner quarters. For live imaging experiments, cells were considered pseudometaphase when they were arrested in prometaphase for 100 minutes or with or without obvious congression defect. Immunofluorescence experiments from Figures 3F-K were processed in ImageJ/FIJI by using the background subtraction feature then measuring intensities. For Figs 3F-3K, the area to be analyzed was specified with the free hand tool (3F, 3G, 3J, 3K: nucleus; 3H, 3I: centrosome). Statistical analysis was performed using Graph Pad9 software using Chi squared, Anova, and unpaired t-test as indicated in the figure legends. Error bars represent the standard error of the mean.

## KEY RESOURCES TABLE

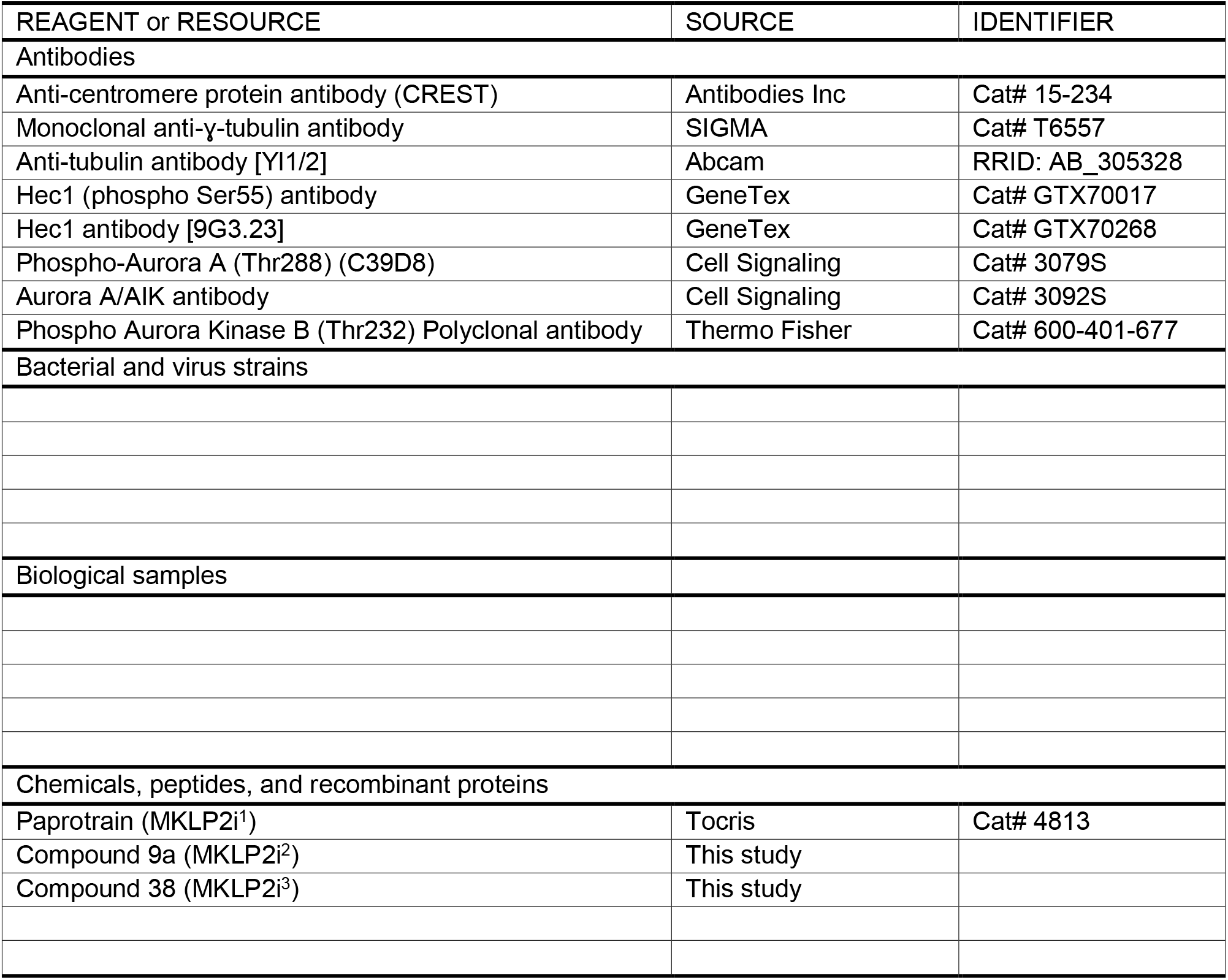
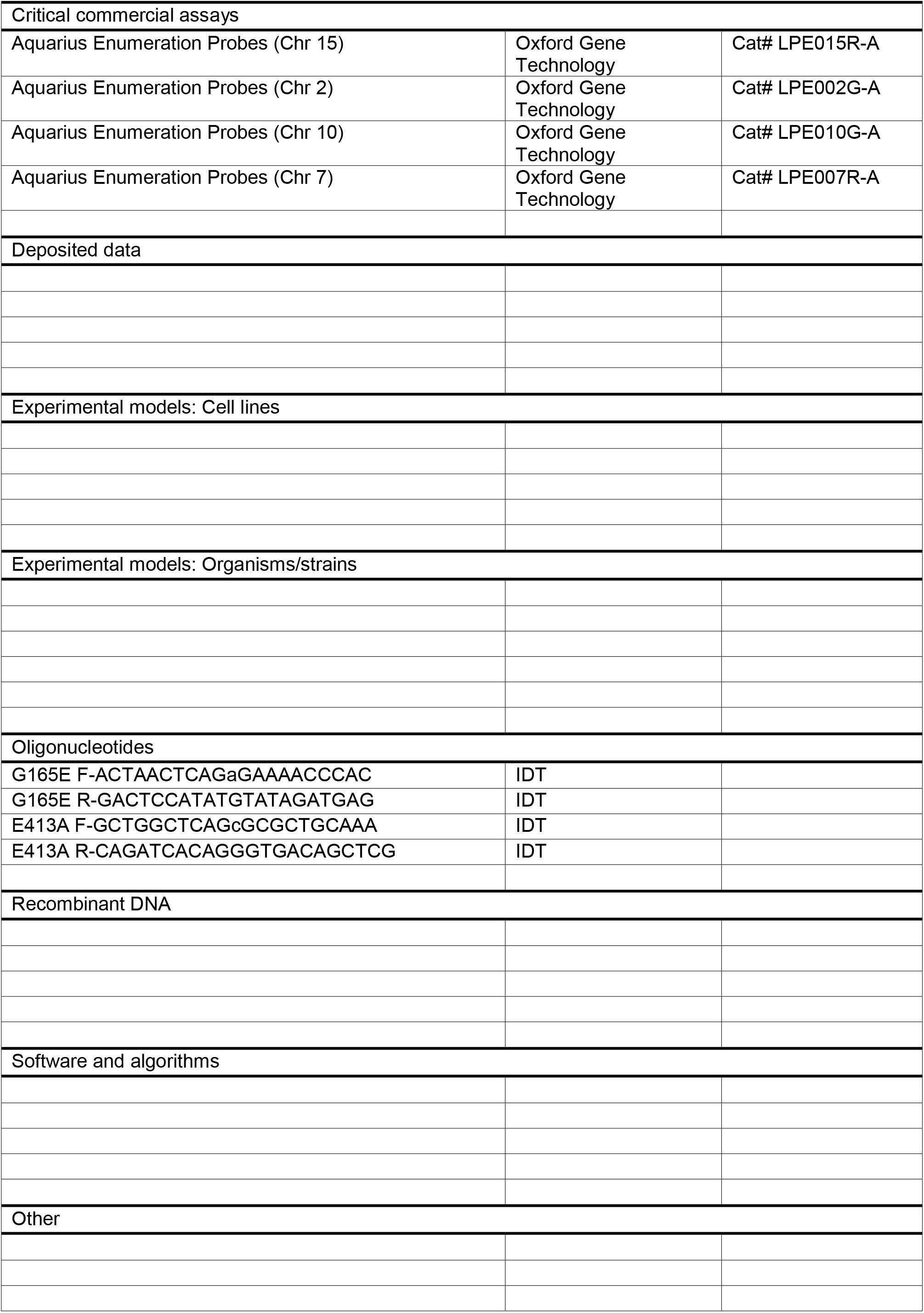
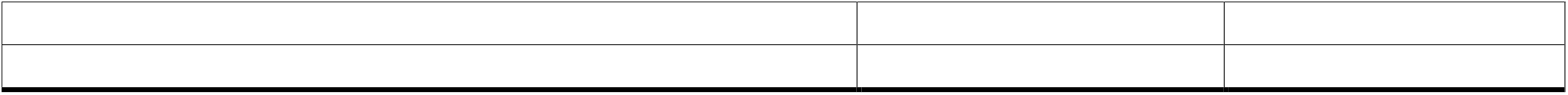

## References

Adriaans, I. E., Hooikaas, P. J., Aher, A., Vromans, M. J. M., Van Es, R. M., Grigoriev, I., Akhmanova, A. & Lens, S. M. A. 2020. MKLP2 Is a Motile Kinesin that Transports the Chromosomal Passenger Complex during Anaphase. Curr Biol, 30, 2628-2637.e9.

Ben-David, U. & Amon, A. 2020. Context is everything: aneuploidy in cancer. Nature Reviews Genetics, 21, 44–62.

Browning, H., Hackney, D. D. & Nurse, P. 2003. Targeted movement of cell end factors in fission yeast. Nat Cell Biol, 5, 812–8.

Carter, S. L., Eklund, A. C., Kohane, I. S., Harris, L. N. & Szallasi, Z. 2006. A signature of chromosomal instability inferred from gene expression profiles predicts clinical outcome in multiple human cancers. Nature Genetics, 38, 1043–1048.

Chmátal, L., Yang, K., Schultz Richard M. & Lampson Michael A. 2015. Spatial Regulation of Kinetochore Microtubule Attachments by Destabilization at Spindle Poles in Meiosis I. Current Biology, 25, 1835–1841.

Deluca, J. G. 2017. Aurora A Kinase Function at Kinetochores. Cold Spring Harb Symp Quant Biol, 82, 91–99.

Deluca, K. F., Herman, J. A. & Deluca, J. G. 2016. Measuring Kinetochore-Microtubule Attachment Stability in Cultured Cells. Methods Mol Biol, 1413, 147–68.

Deluca, K. F., Meppelink, A., Broad, A. J., Mick, J. E., Peersen, O. B., Pektas, S., Lens, S. M. A. & Deluca, J. G. 2018. Aurora A kinase phosphorylates Hec1 to regulate metaphase kinetochore-microtubule dynamics. J Cell Biol, 217, 163–177.

Ertych, N., Stolz, A., Valerius, O., Braus, G. H. & Bastians, H. 2016. Ȗlt;emȖgt;CHK2Ȗlt;/emȖgt;– Ȗlt;emȖgt;BRCA1Ȗlt;/emȖgt; tumor-suppressor axis restrains oncogenic Aurora-A kinase to ensure proper mitotic microtubule assembly. Proceedings of the National Academy of Sciences, 113, 1817.

Ferrero, H., Corachán, A., Quiñonero, A., Bougeret, C., Pouletty, P., Pellicer, A. & Domënguez, F. 2019. Inhibition of KIF20A by BKS0349 reduces endometriotic lesions in a xenograft mouse model. Mol Hum Reprod, 25, 562–571.

Gurden, M. D., Anderhub, S. J., Faisal, A. & Linardopoulos, S. 2016. Aurora B prevents premature removal of spindle assembly checkpoint proteins from the kinetochore: A key role for Aurora B in mitosis. Oncotarget, 9, 19525–19542.

Hill, S. J., Rolland, T., Adelmant, G., Xia, X., Owen, M. S., Dricot, A., Zack, T. I., Sahni, N., Jacob, Y., Hao, T., Mckinney, K. M., Clark, A. P., Reyon, D., Tsai, S. Q., Joung, J. K., Beroukhim, R., Marto, J. A., Vidal, M., Gaudet, S., Hill, D. E. & Livingston, D. M. 2014. Systematic screening reveals a role for BRCA1 in the response to transcription-associated DNA damage. Genes & development, 28, 1957–1975.

Hümmer, S. & Mayer, T. U. 2009. Cdk1 negatively regulates midzone localization of the mitotic kinesin Mklp2 and the chromosomal passenger complex. Curr Biol, 19, 607–12.

Kapoor, T. M., Lampson, M. A., Hergert, P., Cameron, L., Cimini, D., Salmon, E. D., Mcewen, B. F. & Khodjakov, A. 2006. Chromosomes can congress to the metaphase plate before biorientation. Science (New York, N.Y.), 311, 388–391.

Katayama, H., Sasai, K., Kloc, M., Brinkley, B. R. & Sen, S. 2008. Aurora kinase-A regulates kinetochore/chromatin associated microtubule assembly in human cells. Cell Cycle, 7, 2691–2704.

Kim, Y., Holland, A. J., Lan, W. & Cleveland, D. W. 2010. Aurora Kinases and Protein Phosphatase 1 Mediate Chromosome Congression through Regulation of CENP-E. Cell, 142, 444–455.

Kitagawa, M., Fung, S. Y., Hameed, U. F., Goto, H., Inagaki, M. & Lee, S. H. 2014. Cdk1 coordinates timely activation of MKlp2 kinesin with relocation of the chromosome passenger complex for cytokinesis. Cell Rep, 7, 166–79.

Krenn, V. & Musacchio, A. 2015. The Aurora B Kinase in Chromosome Bi-Orientation and Spindle Checkpoint Signaling. Frontiers in Oncology, 5.

Krupina, K., Kleiss, C., Metzger, T., Fournane, S., Schmucker, S., Hofmann, K., Fischer, B., Paul, N., Porter, I. M., Raffelsberger, W., Poch, O., Swedlow, J. R., Brino, L. & Sumara, I. 2016. Ubiquitin Receptor Protein UBASH3B Drives Aurora B Recruitment to Mitotic Microtubules. Dev Cell, 36, 63–78.

Labrière, C., Talapatra, S. K., Thoret, S., Bougeret, C., Kozielski, F. & Guillou, C. 2016. New MKLP-2 inhibitors in the paprotrain series: Design, synthesis and biological evaluations. Bioorganic & Medicinal Chemistry, 24, 721–734.

Lampson, M. A. & Cheeseman, I. M. 2011. Sensing centromere tension: Aurora B and the regulation of kinetochore function. Trends in cell biology, 21, 133–140.

Lampson, M. A., Renduchitala, K., Khodjakov, A. & Kapoor, T. M. 2004. Correcting improper chromosome-spindle attachments during cell division. Nat Cell Biol, 6, 232–7.

Landino, J., Norris, S. R., Li, M., Ballister, E. R., Lampson, M. A. & Ohi, R. 2017. Two mechanisms coordinate the recruitment of the chromosomal passenger complex to the plane of cell division. Mol Biol Cell, 28, 3634–3646.

Pouletty, P. 2019. New derivatives of indole for the treatment of endometriosis. United States patent application 16 / 344, 390

Serena, M., Bastos, R. N., Elliott, P. R. & Barr, F. A. 2020. Molecular basis of MKLP2-dependent Aurora B transport from chromatin to the anaphase central spindle. The Journal of cell biology, 219, e201910059.

Shihavuddin, A., Basu, S., Rexhepaj, E., Delestro, F., Menezes, N., Sigoillot, S. M., Del Nery, E., Selimi, F., Spassky, N. & Genovesio, A. 2017. Smooth 2D manifold extraction from 3D image stack. Nature Communications, 8, 15554.

Stumpff, J., Wagenbach, M., Franck, A., Asbury, C. L. & Wordeman, L. 2012. Kif18A and chromokinesins confine centromere movements via microtubule growth suppression and spatial control of kinetochore tension. Dev Cell, 22, 1017–29.

Tcherniuk, S., Skoufias, D. A., Labriere, C., Rath, O., Gueritte, F., Guillou, C. & Kozielski, F. 2010. Relocation of Aurora B and Survivin from Centromeres to the Central Spindle Impaired by a Kinesin-Specific MKLP-2 Inhibitor. Angewandte Chemie International Edition, 49, 8228–8231.

Wimbish, R. T. & Deluca, J. G. 2020. Hec1/Ndc80 Tail Domain Function at the Kinetochore-Microtubule Interface. Frontiers in cell and developmental biology, 8, 43–43.

Ye Anna A., Deretic, J., Hoel Christopher M., Hinman Albert W., Cimini, D., Welburn Julie P. & Maresca Thomas J. 2015. Aurora A Kinase Contributes to a Pole-Based Error Correction Pathway. Current Biology, 25, 1842–1851.

