## Supplementary figures and images for "MKLP2 functions in early mitosis to ensure proper chromosome congression"

### Suppl Fig 1

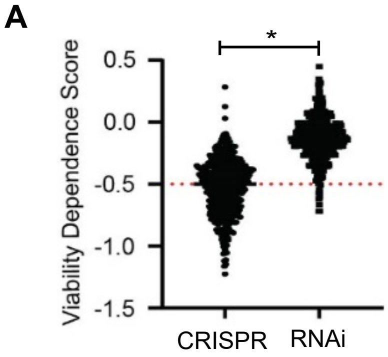


Supplemental Figure 1
